# Evolutionary dynamics of a lethal recessive allele in reintroduced fragmented lynx populations

**DOI:** 10.1101/2025.11.06.686959

**Authors:** Janna Niehaus, Michelle Imlau, Saskia Keller, Iris Marti, Christine Breitenmoser, Anna Letko, Vidhya Jagannathan, Christine Grossen, Tosso Leeb, Mirjam Pewsner

## Abstract

Conservation programs worldwide use reintroductions and translocations as management tools to support endangered species. Such efforts often face genetic challenges, as small populations and restricted gene flow can lead to inbreeding and loss of genetic diversity. Inbreeding depression is a well-known risk in small, isolated populations, where reduced genetic diversity can compromise fitness and threaten population persistence. While correlations between inbreeding and reduced fitness are well documented, so far, no causal variant for a fitness-relevant phenotype has been described in a wild population. Here, we combine long-term health and population data with genomic analyses to identify a recessive lethal allele responsible for extensive organ mineralization in young Eurasian lynx (*Lynx lynx*) from a reintroduced population in Switzerland. The causal variant is located in *FGF23* and predicted to impair secretion of fibroblast growth factor 23, an important regulator of phosphate homeostasis. The deleterious allele was likely introduced from one of the lynx source populations and subsequently increased in frequency through founder effects and inbreeding. All three Swiss lynx populations show high inbreeding levels (FROH > 0.4), reflecting substantial loss of genetic diversity. Our findings uniquely demonstrate how demographic history, founder effects, genetic drift, and inbreeding interact to shape the fate of a recessive deleterious allele, highlighting the importance of multiscale monitoring in wildlife conservation.

## Main

Every species reintroduction is not only an ecological intervention, but also an inadvertent experiment in evolutionary genetics. Reintroductions are a widely used conservation strategy to prevent extinction, yet they carry important genetic risks. Bottlenecks and small founder populations not only lead to loss of genetic diversity and thereby loss of adaptive potential, they can also accelerate the accumulation of harmful mutations, posing an additional risk to fitness and threatening long-term viability. The role of deleterious mutations for survival of wild populations and how this should affect management decisions of endangered species is a central question for ongoing research. Recent studies have evaluated genome-wide estimates of putatively harmful mutations as a consequence of small population size (e.g. Dussex et al., 2021; Grossen et al., 2020; Xue et al., 2015, reviewed in Dussex et al. 2023). Yet while population genetic models consistently predict substantial risks from such mutations (Charlesworth & Willis, 2009), empirical examples from wild populations linking deleterious mutations with fitness declines remain scarce (Hasselgren et al., 2024; Kardos et al., 2023; Stoffel et al., 2021a). Inbreeding depression, which mainly arises through expression of harmful alleles, has been associated with reduced reproductive output, lower juvenile survival, compromised immunity, shortened lifespans and population viability (Hasselgren et al., 2024; Kardos et al., 2023; Stoffel et al., 2021b; Bozzuto et al., 2019; Huisman et al., 2016; Reid et al., 2003; reviewed in Keller & Waller, 2002). However, few studies bridge the gap between specific genomic evidence and phenotypic consequences. In a wild population of the primitive domestic Soay sheep, variation within *RXFP2* was shown to be linked to horn growth and survival resulting in heterozygote advantage (Johnston et al., 2013). In the small and isolated Similipal tiger population an *LVRN* missense variant was identified as the basis of a pseudomelanistic phenotype (Sagar et al., 2021). Although no fitness effects have been detected, the tiger example illustrates how genetic drift can lead to the spread of novel traits in inbred populations. Such findings highlight the importance of monitoring genetic processes in small, isolated, and reintroduced populations. Yet up to date, no causal variant for a specific fitness-relevant or even lethal phenotype has been described in a wild population.

The reintroduction of the Eurasian lynx (*Lynx lynx*) in Switzerland provides a unique opportunity to study the long-term genetic consequences of conservation management actions. Once extirpated from central Europe, the species was reestablished in the 1970s through the translocation of about 30 lynx from the Carpathians into the Swiss Alps and Jura Mountains (KORA, 2022; Breitenmoser, 1998; Fig. 1). Since then, the total number of Swiss lynx has grown to roughly 360 individuals (Vogt et al., 2025) and has been continuously monitored for over 50 years. Despite this increase in population size, genetic diversity has been severely reduced (KORA, 2022). Thirty years after the reintroduction, the Alpine and Jura populations remained largely isolated. To improve connectivity and enhance the lynx distribution, 12 animals were translocated to northeastern Switzerland (NE-CH) between 2001 and 2008 (Fig. 1). Of these, only seven individuals were confirmed to have contributed to the establishment of the new population, and gene flow among the three regions remains limited, resulting in persistently low genetic diversity (Mueller et al., 2022).

**Figure 1.**
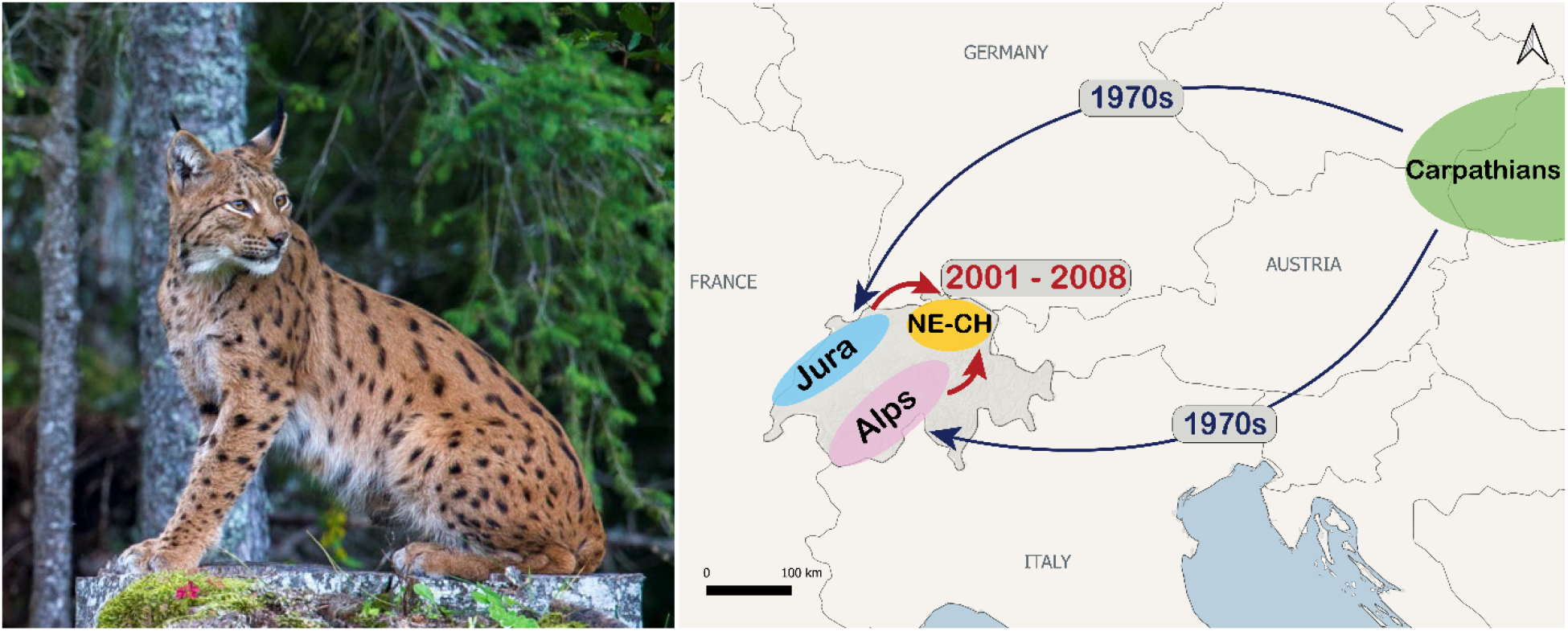
The Eurasian lynx (Lynx lynx) in Switzerland. Left: female lynx in the Jura mountains. Right: history and routes of lynx reintroductions and translocations in Switzerland.

Alongside demographic and genetic monitoring, Switzerland has implemented an intensive health surveillance based on captures and clinical examinations of live lynx as well as postmortem analyses of all lynx found dead. This has revealed concerning health conditions in recent years, including recurrent cases of heart disease (Ryser-Degiorgis et al., 2020) and severe soft tissue mineralizations in young lynx. Both are suspected to have a genetic basis. Here, we focus on the mineralization phenotype and describe how reintroduction history, followed by low genetic diversity led to the expression of a lethal recessive allele in the Eurasian lynx.

## Results

### Regional Clustering of Fatal Multi-Organ Mineralization in Young Eurasian Lynx

The Eurasian lynx is a protected species in Switzerland and closely monitored. Post-mortem examinations of all dead lynx (found dead or culled due to health issues or repeated livestock predation) are carried out at the Institute for Fish- and Wildlife Health (FIWI). Since the 1980s biological samples are routinely bio-banked and since 2000 findings are systematically recorded in a database compiling health data of all lynx examined. To investigate the occurrence of the mineralization phenotype in more detail, a retrospective diagnostic case review, including histological reassessment, was performed. This revealed nine animals (six found dead and three weak and emaciated) sharing a highly similar histopathological presentation in multiple organs (Suppl. Table S1). All affected individuals were young and of equal sex-ratio (Marti & Ryser-Degiorgis, 2018). All were found in the northeastern part of Switzerland (NE-CH). On histopathology, the kidneys, lungs, and stomach consistently exhibited comparable patterns of extensive mineralization. In the kidneys, the renal cortex showed diffuse mineralization (Fig. 2A, D). All cases also showed widespread pulmonary mineralization (Fig. 2B, E). In eight animals, marked mineralization occurred in the stomach, predominantly within the mucosal layer (Fig. 2C, F). Severe autolysis limited histopathological assessment of the stomach in one animal. The severity and distribution of mineralization in all cases were incompatible with normal organ function and were considered ultimately fatal. A detailed overview of the pathological findings and case background is compiled in the Supplementary Material (Suppl. Table S1).

**Figure 2.**
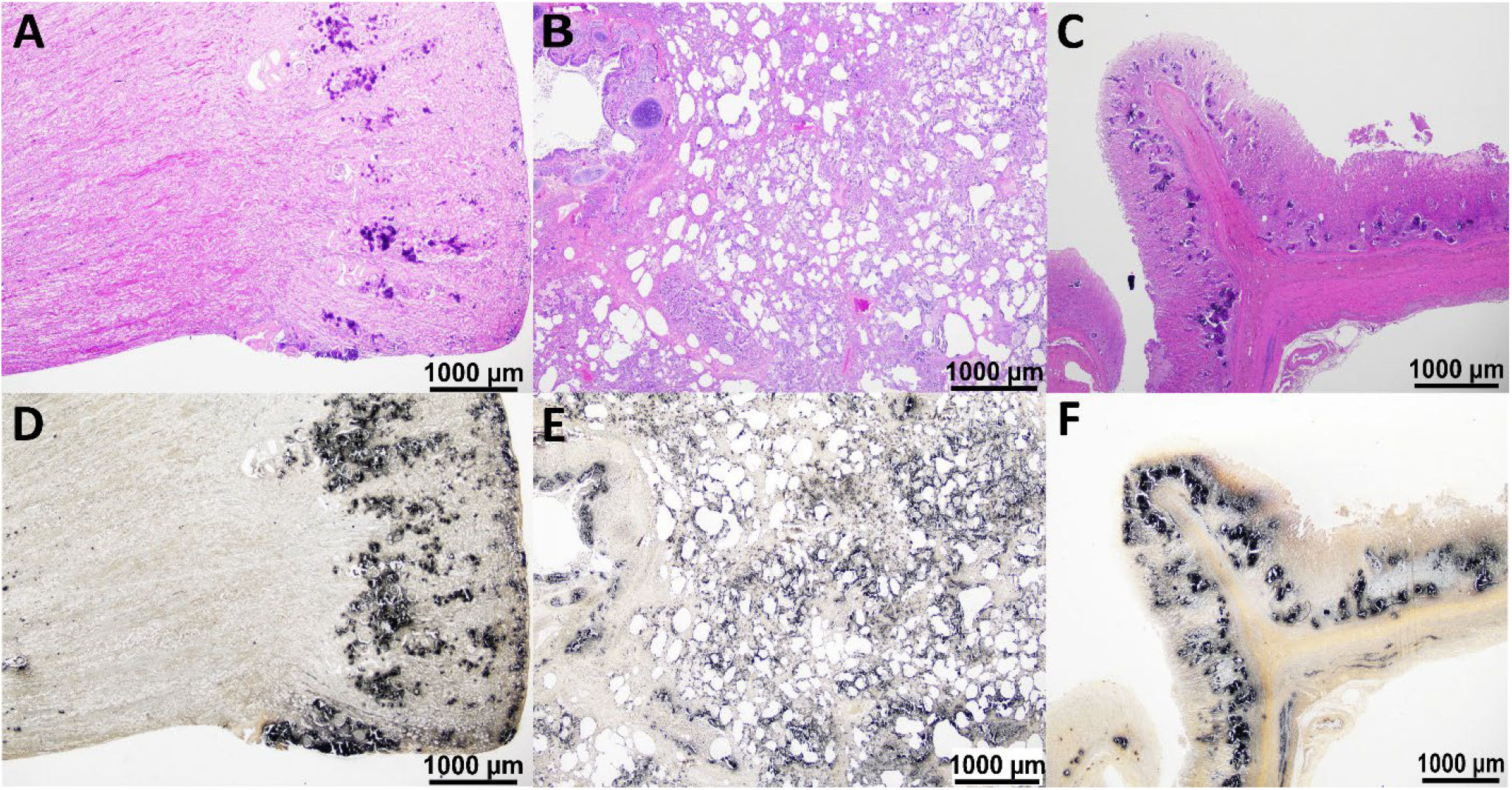
Histopathological findings in Eurasian lynx (Lynx lynx) affected with metastatic mineralizations. Top row showing affected kidney (A), lung (B), and stomach (C) in H&E stain; Mineralization is visible as dark purple, crystalline structures. Bottom (D-F) shows same organ sections as in top row, in von Kossa stain, highlighting mineralized areas in black.

### Strong Population Differentiation and High Levels of Inbreeding

Given the reintroduction history of the affected NE-CH population which experienced two severe bottlenecks (Fig. 1), the observed pathology was hypothesized to result from a deleterious recessive allele and inbreeding. To test this hypothesis and search for a possible genetic basis of the observed mineralization phenotype, we generated whole-genome sequencing (WGS) data of 50 lynx individuals comprising nine mineralization-cases from NE-CH and 41 controls, representing all three Swiss lynx populations (Alps = 20, Jura = 13, NE-CH = 8; Suppl. Fig. S1, Suppl. Table S2). The WGS dataset contained 1,899,640 high-quality bi-allelic autosomal variants (Suppl. Table S3). A principal component analysis (PCA) based on the genotypes at this subset of markers revealed clear genetic structure among the three populations (Fig. 3A, Suppl. Fig. S2). All nine affected individuals clustered within the NE-CH population with no separation of cases from controls (Fig. 3B). The strong among-population differentiation was confirmed by the STRUCTURE-like analysis (Suppl. Fig. S3).

**Figure 3.**
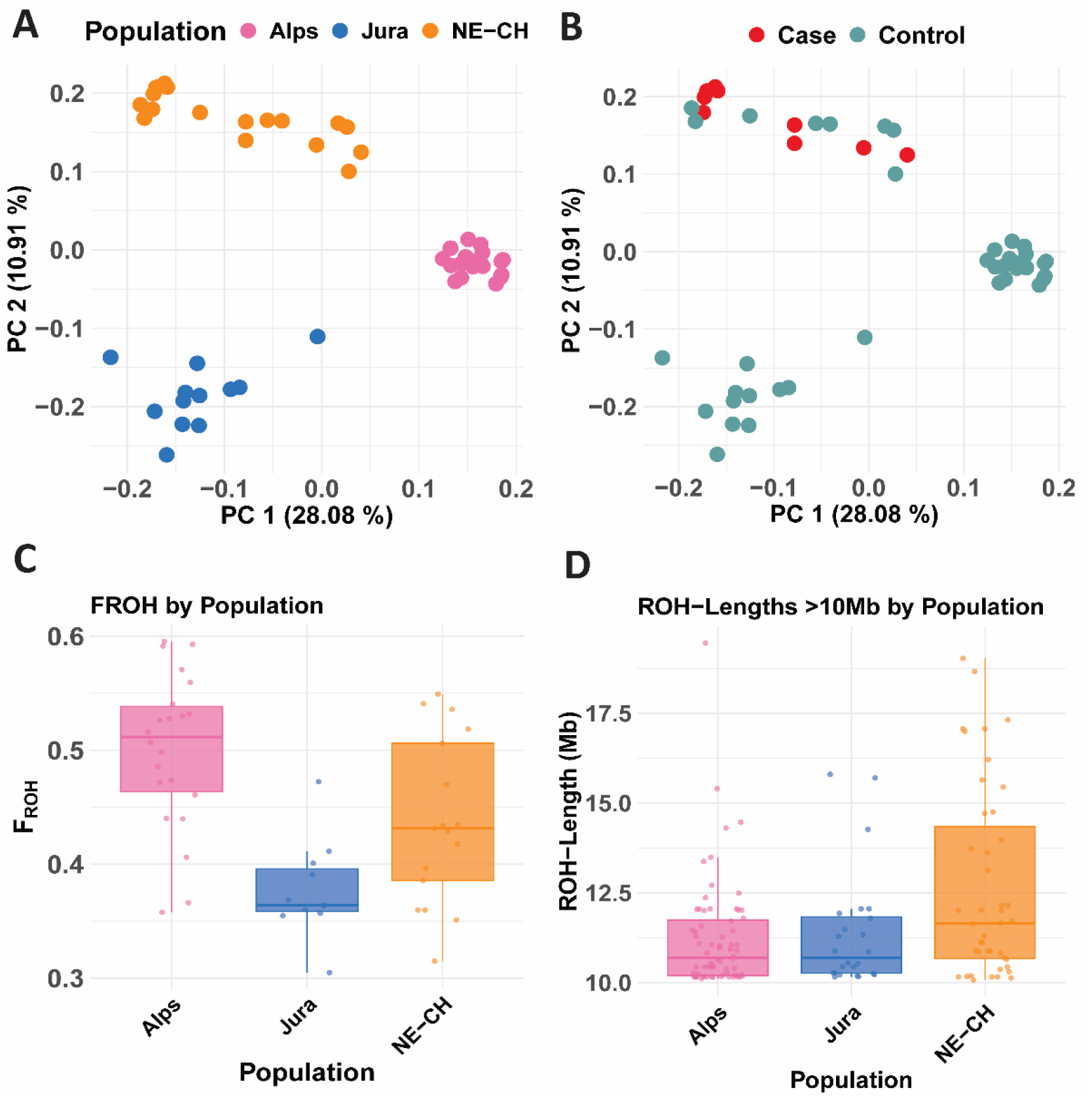
Principal component and inbreeding analyses reveal strong population structure and high inbreeding levels. (A) Principal Component Analysis (PCA) plot of the first two principal components (PCA1 vs. PCA2), demonstrating clear genetic separation among all three Swiss lynx populations. (B) PCA plot coloured by phenotypic group, red dots indicate the nine mineralization cases. (C) Boxplots of individual FROH values (proportion of the genome in ROHs >1 Mb) for each population, highlighting high inbreeding levels across all populations. (D) Distribution of ROH lengths >10Mb across the three populations.

Runs of homozygosity (ROH) analysis across the autosomal genome confirmed very high levels of inbreeding in all three lynx populations, consistent with mating among siblings and between parents and offspring (Fig. 3C, D). The Alpine population exhibited the highest levels of inbreeding, the Jura population showed the lowest, while the NE-CH population displayed intermediate levels and a wider range of FROH values (proportion of genome within ROH larger than 1 Mb), reflecting heterogeneous inbreeding patterns (Fig. 3C, more details in Suppl. Fig. S4). Analyses of particularly long ROH segments (> 10 Mb, indicative of recent inbreeding) showed the highest levels in the NE-CH population, consistent with its more recent reintroduction history (Fig. 3D).

### Identification of the Causal Variant for the Lethal Mineralization Phenotype

A genome-wide association study (GWAS) for the mineralization locus, using a further reduced dataset comprising 1,699,701 high-quality markers yielded the strongest association signal (*P* = 4.88 × 10^-8^) on scaffold NW_025814889.1 at position 39,312,590 (Fig. 4). All 9 cases were homozygous at the best associated marker confirming the hypothesized autosomal recessive inheritance.

**Figure 4.**
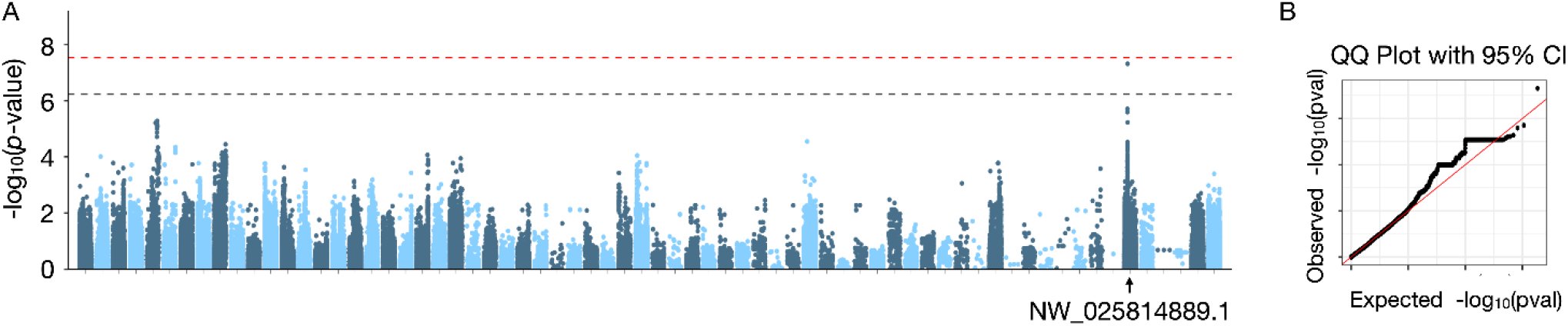
GWAS analysis identified one variant exceeding the suggestive significant threshold. (A) Manhattan plot visualizing the association between genetic variants and the phenotypic trait. The x-axis represents genome positions grouped by scaffolds, and the y-axis indicates the −log_10_(p-value) for each variant. Genome-wide significance and suggestive significance thresholds were calculated using the Bonferroni correction for multiple testing, based on 1,699,701 independent variants tested (P = 0.05/1,699,701 = 2.93 × 10_−8_ for genome-wide significance, marked by red dotted line; and P = 1.0/1,699,701 = 5.88 × 10_−7_ for suggestive significance, marked by black dotted line). One variant exceeded the suggestive significance threshold (P = 4.88 × 10^−7^) on scaffold NW_025814889.1. (B) QQ plot comparing the observed distribution of −log10(p-value) with the expected distribution under the null hypothesis (no association, red line).

Filtering of all 17,328,023 raw variants (Suppl. Table S3) for homozygous alternate genotypes exclusively in the nine cases but in none of the controls identified one single position, which exactly corresponded to the best associated variant in the GWAS, NW_025814889.1:g.39,312,590A>G. It was a missense variant affecting the coding sequence of *FGF23*, XM_047065192.1:c.38T>C. The identified variant is predicted to change leucine-13 in the signal peptide of the FGF23 protein into a proline, XP_046921148.1:p.(Leu13Pro).

*FGF23* encodes fibroblast growth factor 23, which is a main regulator of phosphorous homeostasis in the body. In humans, loss of function has been shown to cause hyperphosphatemia, which can lead to hypermineralizations (Garringer et al., 2008). Protein predictions for the XP_046921148.1:p.(Leu13Pro) variant consistently predicted a pathogenic or disease-causing effect, most likely due to defective recognition and cleavage of the signal peptide (for more details see Supplementary Note S2 and Suppl. Fig S5).

To further validate the genotype-phenotype association, a total of 153 individuals sampled between 1990 and early 2025, for which detailed health data was available, were genotyped for the *FGF23*:c.38T>C variant. They included the 50 animals from the WGS experiment. This confirmed a perfect genotype-phenotype association (Table 1), identified one historical mineralization case from 1999 in the Alpine population, and provided genetic confirmation for two new histopathological identified cases in NE-CH from 2024, bringing the total number of detected cases to 12.

**Table 1.** Genotype-phenotype association at the FGF23:c.38T>C variant.

| Phenotype | wt/wt | wt/mut | mut/mut |
| --- | --- | --- | --- |
| Mineralization cases (n = 12)<sup>a</sup> | - | - | 12 |
| Controls (n = 141) | 98 | 43 | - |

<sup>a</sup>These included the 9 initially discovered cases, 2 new cases that were sampled in 2024 during the study and one retrospectively identified historical case sampled in 1999 whose diagnosis had not yet been electronically recorded (details in Table S2).

### Deleterious Allele was Introduced to NE-CH from Alpine Population

The oldest of the 12 identified cases had been found dead in 1999 in the Alpine population, more specifically in the region of the southwestern Swiss Alps (SWSA). In the sampling period from 1990-2000, which pre-dated the translocation of animals to NE-CH, the frequency of the deleterious allele was with 11 carriers out of 13 tested individuals very high in the SWSA region (Fig. 5, Suppl. Table S2). We successfully genotyped 10 of the 12 to the NE-CH translocated individuals. Three of them were carriers, two females and one male, and all originated from the SWSA region. Genotyping failed for two individuals due to insufficient DNA quality. However, pedigree data from KORA (2024) suggests that at least one of these individuals, another female, was most likely heterozygous, as it produced heterozygous offspring with a known wildtype mate. This probable additional carrier also originated from the SWSA region. The frequency of deleterious alleles in the SWSA region declined markedly after the sampling period from 1990 – 2000 to 6 carriers out of 36 genotyped individuals sampled between 2012 and 2025. In contrast, the frequency of the deleterious allele increased in the NE-CH population since its foundation. Between 2012 and 2025, 16 out of 27 unaffected individuals were carriers, which is consistent with the high number of animals homozygous for the mutant allele and hence affected with mineralizations detected in this region. So far, we did not discover the mutant allele in the Jura population (Fig. 5, Suppl. Table S2).

**Figure 5.**
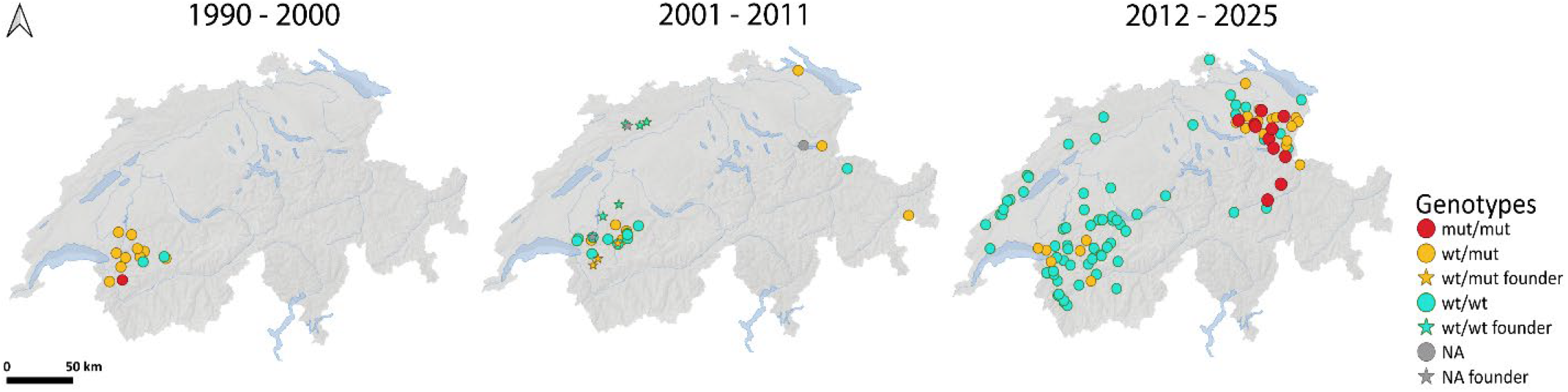
Geographic distribution of lynx genotyped for the FGF23:c.38T>C variant across Switzerland over three time periods. Each map shows the location where individuals were found, with symbols color-coded by genotype. Left map, individuals found in the southwestern Swiss Alps (SWSA) region between 1990 and 2000. Center, individuals from the southwestern Swiss Alps (SWSA) region, all individuals translocated to the north-eastern Switzerland (NE-CH) population (stars) as well as the first individuals born in NE-CH and sampled between 2001 and 2011. Right map, all genotyped individuals sampled between 2012 and 2025.

## Discussion

Combining long-term pathological and population data, conservation genomics, and animal genetics, we identified a recessive allele causing lethal mineralization in young individuals of a recently reintroduced Eurasian lynx population in Switzerland. Genotyping of the founders revealed that the deleterious allele was introduced from the Alpine population, one of the two sources used for the reintroduction (Fig. 1). While already present in the Alpine population for at least 30 years (Fig. 5), the detrimental effect of this allele only became apparent after its introduction to NE-CH through a small number of mostly heterozygous founders and subsequent inbreeding in a small population of less than 25 independent individuals. This highlights the risks posed by genetic bottlenecks and inbreeding and demonstrates the complex interplay of demographic history, selection and restricted gene flow in the emergence, persistence and expression of deleterious alleles.

Levels of inbreeding measured as FROH were well above 0.4 in all three Swiss populations studied here, indicating substantial loss of genetic diversity and high levels of within-group relatedness, consistent with previous reports in Swiss lynx (Breitenmoser & Obexer-Ruff, 2003; Mueller et al., 2022). Comparable FROH values have been observed in the Isle Royale wolf population before translocations saved it from extinction (Robinson et al., 2019). It is widely accepted that high levels of inbreeding pose a threat to population survival through inbreeding depression (Charlesworth & Charlesworth, 1987; Charlesworth & Willis, 2009; Hedrick & Garcia-Dorado, 2016; Kardos et al., 2024). Observations in a number of wild species showed that extensive homozygosity and inbreeding were associated with detrimental phenotypes, such as chondrodystrophy in the California condor (Ralls et al., 2000), reduced reproductive performance in Scandinavian arctic foxes (Hasselgren et al., 2024; Norén et al., 2016), and various morphological and health problems in Florida panthers (Roelke et al., 1993), and wolves from Isle Royale (Robinson et al., 2019). While these examples mostly reported correlations between inbreeding and reduced fitness, we here gained additional insights on the molecular level and demonstrate a direct causal link between a specific recessive allele and mortality.

For an improved understanding of the origin and past dynamics of the lethal variant, we used targeted genotyping to trace the deleterious allele back to at least 1991 in the SWSA region of the Alpine population, one of the sources of the affected NE-CH population (Fig. 5). Interestingly, the allele frequency was locally relatively high in the 1990s, yet only one mineralization case was detected in this population, and no apparent population effects could be observed. Our data suggests that the mutant allele frequency in the SWSA region has since decreased continuously, while still persisting at low levels. This decline may reflect the process of purging, where recessive deleterious alleles are progressively removed when expressed in homozygotes (Glémin, 2003; Kirkpatrick & Jarne, 2000). However, while purging can sometimes reduce genetic load and contribute to population viability, even lethal alleles can remain masked in a heterozygous state and the effectiveness of purging strongly depends on population size, demographic history, and the strength of selection (Robinson et al., 2023; Pérez-Pereira et al., 2022). In the SWSA lynx, the decline in allele frequency was likely influenced by multiple mechanisms. Besides some purging and the removal of heterozygotes through translocation to NE-CH, the random effects of genetic drift but also immigration of lynx from neighboring regions into the newly vacant territories may have further diluted the frequency of the deleterious allele.

Contrary to SWSA, in NE-CH the allele became problematic after the introduction through several heterozygous founders. It is known that some of these founders contributed disproportionately to the gene pool of the newly established population (KORA, 2024). Such unequal founder contributions are a well-known risk in reintroductions (Kivisild, 2013). Additionally, establishment of the deleterious allele was likely accelerated by small effective population size and hence genetic drift, restricted dispersal and demographic stochasticity, increasing the probability of homozygous, diseased individuals. Furthermore, given that individuals from two diverged populations were used for the translocation (in addition to the Alps also individuals from the Jura), masking of deleterious alleles in admixed individuals may have further limited purifying selection during the first few generations. Similar to the case of momentary genetic rescue in the wolf population on Isle Royale, such admixed individuals carrying masked alleles may have profited from increased fitness. However, if a population doesn’t increase in size fast enough and genetic diversity is small (here based on seven founders), inbreeding can quickly become a risk for population survivability (Franklin, 1980). The commonly advised 50/500 rule recommends an effective population size *N*_*e*_ of at least 50 individuals to avoid short-term inbreeding depression and 500 to retain long-term adaptive potential (Franklin, 1980) where *N*_*e*_ is usually smaller than census size, often reported to be around 10-30% (reviewed in Hoban et al., 2020). However, even these values are discussed as potentially insufficient (Frankham et al., 2014). As of winter 2024/2025, the NE-CH population was estimated to include approximately 21 (95% CI: 21–29) independent individuals, with population growth having stagnated after 2016 (Sterrer et al., 2025; KORA, 2024). In such a small and highly inbred population, genetic drift can counteract purging. While homozygous individuals die before reproducing, thereby reducing the frequency of the deleterious allele, the high proportion of heterozygotes with no fitness reduction allows the deleterious allele to persist at moderate frequency and less deleterious alleles may even approach fixation (Robinson et al., 2023). In the isolated Similipal tiger population, drift likely facilitated the rise of the recessive melanistic allele, which apparently has no detrimental effect (Sagar et al., 2021). Although the presence of the deleterious allele and inbreeding in general may have contributed to the observed stagnation of population growth in the NE-CH, other factors could also play a role, making definite conclusions difficult. Initial population growth is often seen in reintroduced species with abundant resources, but growth rates typically slow down as resource and space limitations are reached (Sibly & Hone, 2003). This could mean that the NE-CH population, which experienced significant growth between 2009 and 2015 (KORA, 2024), may have approached its carrying capacity at ∼20-30 individuals. However, comparisons with other Alpine regions show that higher lynx densities are possible (Kunz et al., 2020, 2021; Sterrer et al., 2022). Additionally, human-mediated removal of four females, for a European transboundary translocation project between 2016 and 2020 may have further negatively influenced population growth (KORA, 2024).

The identified variant causing lethal mineralization in Swiss lynx demonstrates how deleterious recessive alleles can persist and remain mostly hidden in a source population, but become problematic after reintroduction, where founder effects and inbreeding take effect. It underscores the importance of establishing reintroduced populations with a sufficiently large and genetically diverse founder base. In Switzerland, with at most 30 individuals founding the Swiss lynx populations and only seven in the NE-CH, numbers were far below the recommended thresholds of the 50/500 rule (Franklin, 1980). Importantly, beyond numbers of founding individuals, long-term viability depends on additional factors including post-introduction population growth and population connectivity to reduce isolation, facilitate gene flow and mitigate inbreeding (Benson et al., 2016; Hedrick & Garcia-Dorado, 2016; Weeks et al., 2011).

This study further emphasizes the need for long-term intensive post-reintroduction monitoring, both genetically and health-related, to ensure early detection of emerging threats to population viability. To provide a basis for informed management of the mineralization phenotype, future research should focus on monitoring allele frequency and demographic trends in the Swiss lynx populations. As all three Swiss lynx populations show high levels of inbreeding, genetic rescue may become an indispensable tool to restore diversity, counteract inbreeding depression and enable adaptive potential for future environmental changes. At the same time conservation strategies need to prioritize the establishment of connectivity and prevention of isolation to ensure gene flow and reduce the risk of introducing other deleterious genetic variants into an inbreeding setting.

To our knowledge, this is the first study in a wild species to demonstrate a specific recessive allele directly causing mortality linked to inbreeding. This example illustrates the theoretically long-known dynamics and risks of inbreeding in small and reintroduced populations. It underscores the importance of large and genetically diverse founder populations, maintaining connectivity, implementing continuous health and demographic monitoring and applying genomic tools to guide management decisions from the start.

## Methods

### Sample Collection, Case Identification and Histopathology

For this study, samples were obtained through the Swiss national health monitoring program of the Eurasian lynx. Within this framework, post-mortem examinations and immobilizations of live lynx are systematically accompanied by standardized sampling procedures. Organ tissues and blood are routinely collected and archived in a biobank. Additional tissue samples are fixed in 4% buffered formalin, processed, and stained with hematoxylin and eosin (H&E) following established protocols of the Institute of Animal Pathology, Vetsuisse Faculty, University of Bern. All H&E-stained histological slides used in this study originated from this standardized sampling and examination protocol. Furthermore, the Institute for Fish and Wildlife Health (FIWI) maintains a lynx database with information and examination results on every lynx examined by the institute. Monitoring started in the 1980s, but systematic electronic record-keeping was only implemented around 2000, resulting in incomplete data for earlier years. This database served as the basis for identifying the cases included in this study. The database was searched for cases with multifocal, moderate, or severe mineralizations, which were then further investigated. The corresponding histological slides were subsequently re-analyzed by two independent reviewers, including one board-certified veterinary pathologist. The analysis focused on the presence and amount of mineralization, fibrosis and inflammation. To prevent bias, the assessment was done blinded. Cases with similar histopathological findings regarding affected organs, mineralized structures, and severity were then selected and analyzed in more detail. For analysis of presence and severity of mineralization, von Kossa staining (Kiernan, 2015) was performed on all slides of the investigated animals. To examine fibrosis, we used either elastica-van-Gieson, van-Gieson or Masson-trichrome staining (Kiernan, 2015). Based on the histopathological findings, a case group of nine animals was identified. For further genetic analyses a control group comprising 41 individuals was selected. Those individuals showed no evidence of abnormal mineralization or kidney dysfunction based on their pathological findings or clinical and laboratory examinations. The control samples included lynx from all three Swiss populations (Fig. 1). All 50 individuals assigned to the case/control groups were subjected to whole-genome sequencing.

### DNA Extraction, Whole-Genome Sequencing and Bioinformatic Processing

For whole-genome sequencing and targeted genotyping, DNA was extracted from 153 tissue or EDTA blood samples using the Maxwell Tissue DNA Kit or the Maxwell RSC Whole Blood DNA Kit, respectively (both kits from Promega, Dübendorf, Switzerland) following the manufacturer’s protocol. Samples were taken from the biobanks of FIWI (n=142) and the Foundation for Carnivore Ecology and Wildlife Management (KORA, n=11).

Whole-genome sequencing of 50 individuals was performed at the Next Generation Sequencing Platform of the University of Bern. Briefly, PCR-free DNA libraries with an insert size of approximately 500 bp were prepared. Libraries were individually indexed, pooled and sequenced with 2 × 150 bp on an S4 flow cell of an Illumina NovaSeq 6000 Instrument (Illumina, San Diego, CA, USA). The reads were cleaned with fastp (v0.23.4). The cleaned fastq data was mapped to the Bobcat (*L. rufus*), mLynRuf1.p (NCBI RefSeq GCF_022079265.1) genome reference assembly using BWA-mem2 (ver 2.2.1) with the parameters “K 100000000 –Y”. The aligned file was duplicate marked and processed according to the GATK best practices workflow (https://gatk.broadinstitute.org/hc/en-us/articles/360035535932-Germline-short-variant-discovery-SNPs-Indels). The generated vcf file was hard filtered for single nucleotide variants (SNVs) using the following parameters: QD < 2.0, FS > 60, SOR > 4.0, MQ < 30.0, MQRankSum < -5.0; MQRankSum > 5.0, ReadPosRankSum < -8.0, ReadPosRankSum. > 8.0. The variants were annotated using SnpEff with the genome reference annotation version GCF_022079265.1-RS_2023_03. Sequence data were deposited in the European Nucleotide Archive; accession numbers and additional details are provided in Supplementary Table S2. The dataset was further processed using VCFtools (v0.1.16, GCC 10.3.0), retaining only biallelic SNVs, with genotype quality (GQ) ≥ 20, missingness rate ≤ 0.2 and a minor allele frequency (MAF) ≥ 0.05. To identify sex-linked scaffolds, scaffolds from the *L. rufus* genome were aligned to the chromosome-level genome assembly of the Canadian lynx (*L. canadensis)* (NCBI RefSeq GCF_007474595.2) using the online tool D-GENIES (v.1.5.0). Scaffolds aligning to the sex chromosomes of *L. canadensis* were classified as sex-linked in *L. rufus* and excluded from downstream analyses. As additional validation, comparison of the sequencing coverage between male and female samples, expecting differences consistent with sex-linked regions, was carried out.

### Population Genetic Analyses: Population Structure and Runs of Homozygosity

A principal component analysis (PCA) was performed using PLINK (--pca, v.1.9) and visualized in R (v. 4.4.1.). Additionally, to account for population structure and cryptic relatedness a centered genetic relatedness matrix (-gk 1) was calculated in GEMMA (v.0.98.5).

To assess population structure in large genomic datasets, we used the R package *LEA* (v.4.4.1; Frichot & François, 2015), which implements a STRUCTURE-like approach based on sparse non-negative matrix factorization (sNMF). The filtered VCF file with 1,899,640 variants was converted to .geno format using the vcf2geno() function. Ancestry coefficients were estimated using the snmf() function for K values ranging from 2 to 5, with 10 replicates per K. The cross-entropy criterion was used to evaluate model fit, and an entropy plot was generated to identify the optimal number of ancestral populations. For visualization, the estimated ancestry coefficients of the best runs were merged with sample metadata and plotted using ggplot2, showing individual ancestry proportions grouped by the three populations.

To ensure higher reliability of variant sites, the dataset was filtered using a stricter quality-by-depth (QD) threshold of >15 prior to a runs of homozygosity (ROH) analysis (1,785,573 variants; results based on the less stringent filter with QD >2 are shown in Supplementary Fig. 4). ROH detection was performed with BCFtools (v.1.12, GCC 10.3.0). Given the small sample sizes per population, the default alternative allele frequency (--AF-dflt) was set to 0.4, and a genotype quality filter of -G30 was applied to exclude low-confidence genotype calls. ROHs were visualized in R using the *dplyr* and *ggplot2* packages. The inbreeding coefficient based on ROH (FROH) was calculated as the total length of ROH segments longer than 1 Mb divided by the autosomal genome length, which was determined by summing scaffold lengths after excluding the previously identified sex-linked scaffolds.

### Genome-wide Association Study (GWAS)

A genome-wide association study (GWAS) was conducted using the 50 animals with whole-genome data (9 cases, 41 controls) and a filtered dataset of 1,699,701 variants (Suppl. Table S3). To correct for relatedness and population structure, we applied GEMMA (v.0.98.5) to compute the genetic kinship matrix and perform linear mixed model analysis prior to the GWAS. A Bonferroni correction was used to calculate the genome-wide significance threshold (P = 0.05/1,699,701 = 2.93 × 10−8), with a suggestive significance threshold set at 1.0/1,699,701 = 5.88 × 10−7. Manhattan and quantile-quantile (QQ) plots were generated using the qqmanh package in R (v.4.4.1).

### Identification of the candidate causative variant

Variant filtering of the complete WGS dataset was performed using a strict filtering approach that required case genotypes to be homozygous alternative and control genotypes to be homozygous reference, heterozygous or having missing genotypes. No regional restrictions or additional selection criteria were applied.

### In silico analyses of the FGF23:p.L13P variant

The *L. rufus* FGF23 protein sequence (NCBI accession: XP_046921148.1) was compared to the orthologous FGF23 sequences from *L. canadensis (*XP_030177644.1) and domestic cat (*Felis catus*, XP_011282059.2). The comparison was performed using BLASTp (version 2.16.0, accessed 2024-11-05) with default parameters.

To predict the pathogenicity of the identified p.L13P missense variant, the tools SNPs&GO (Capriotti et al., 2013) and MutPred2.0 (Pejaver et al., 2020) were applied using the *L. rufus*

FGF23 reference protein sequence. Additionally, the signal peptide characteristics caused by the L13P substitution were assessed with SignalP 6.0 (Teufel et al., 2022).

### Population Screening for the mutant *FGF23* Allele

Genotyping of the XM_047065192.1:c.38T>C variant was performed via direct Sanger sequencing of PCR amplicons. Primers with the sequences F: ATGGCCGTCCTTGTGTATCT and R: AGACCAGGGAGTCAGGGAGT were designed to amplify a 229 bp product (Details in Supplementary Table S4). PCR products were enzymatically cleaned with exonuclease I and shrimp alkaline phosphatase and sequencing was carried out with the forward PCR primer using BigDye Terminator v3.1 Cycle Sequencing Kit (Thermo Fisher). Sequencing products were precipitated using EDTA/ethanol, resuspended in Hi-Di Formamide (Thermo Fisher Scientific), and analyzed on an ABI 3730 DNA Analyzer (Thermo Fisher Scientific). Sequences were analyzed using Sequencher v5.1 software (GeneCodes, Ann Arbor, MI, USA).

## Acknowledgements

We acknowledge the valuable cooperation of the cantonal hunting authorities of Switzerland. Fieldwork and monitoring of the Swiss lynx were conducted in collaboration with the foundation Carnivore Ecology and Wildlife Management (KORA, Ittigen, Switzerland). The health monitoring of lynx was financially supported by the Federal Office for the Environment and the Federal Food Safety and Veterinary Office. We are grateful to our colleagues at the Institute for Fish and Wildlife Health (FIWI, Vetsuisse Faculty, University of Bern, Switzerland) who performed the necropsies and documented the findings, and to Laurent Geslin for providing the photograph used in this paper. This work was supported by the dedicated staff of the Vetsuisse Faculty, University of Bern. We thank the Next-Generation Sequencing Platform of the University of Bern for performing the high-throughput sequencing experiments and the Interfaculty Bioinformatics Unit of the University of Bern for providing high-performance computing infrastructure. The AI language models ChatGPT and DeepL were used to improve the English language and readability of this work.

## References

Benson, J. F., Mahoney, P. J., Sikich, J. A., Serieys, L. E. K., Pollinger, J. P., Ernest, H. B., & Riley, S. P. D. (2016). Interactions between demography, genetics, and landscape connectivity increase extinction probability for a small population of large carnivores in a major metropolitan area. Proceedings of the Royal Society B: Biological Sciences, 283(1837), 20160957. 10.1098/rspb.2016.0957

Bozzuto, C., Biebach, I., Muff, S., Ives, A. R., & Keller, L. F. (2019). Inbreeding reduces long-term growth of Alpine ibex populations. Nature Ecology & Evolution, 3(9), 1359–1364. 10.1038/s41559-019-0968-1

Breitenmoser, C., & Obexer-Ruff, G. (2003). Population and conservation genetics of two re-introduced lynx (Lynx lynx) populations in Switzerland–a molecular evaluation 30 years after translocation. Proceedings of the 2nd Conference on the Status and Conservation of the Alpine Lynx Population (SCALP), 7–9.

Breitenmoser, U., Ch, B., & Capt, S. (1998). Re-introduction and present status of the lynx (Lynx Lynx) in Switzerland. Hystrix : The Italian Journal of Mammalogy. 10.4404/hystrix-10.1-4118

Capriotti, E., Calabrese, R., Fariselli, P., Martelli, P. L., Altman, R. B., & Casadio, R. (2013). WS-SNPs&GO: a web server for predicting the deleterious effect of human protein variants using functional annotation. BMC Genomics, 14 Suppl 3(Suppl 3), S6. 10.1186/1471-2164-14-S3-S6

Charlesworth, D., & Charlesworth, B. (1987). Inbreeding depression and its evolutionary consequences. Annual Review of Ecology and Systematics, 18(1), 237–268. 10.1146/annurev.es.18.110187.001321

Charlesworth, D., & Willis, J. H. (2009). The genetics of inbreeding depression. Nature Reviews Genetics, 10(11), 783–796. 10.1038/nrg2664

Dussex, N., Morales, H. E., Grossen, C., Dalén, L., & van Oosterhout, C. (2023). Purging and accumulation of genetic load in conservation. Trends in Ecology & Evolution, 38(10), 961–969. 10.1016/j.tree.2023.05.008

Dussex, N., van der Valk, T., Morales, H. E., Wheat, C. W., Díez-del-Molino, D., von Seth, J., Foster, Y., Kutschera, V. E., Guschanski, K., Rhie, A., Phillippy, A. M., Korlach, J., Howe, K., Chow, W., Pelan, S., Mendes Damas, J. D., Lewin, H. A., Hastie, A. R., Formenti, G., … Dalén, L. (2021). Population genomics of the critically endangered kākāpō. Cell Genomics, 1(1), 100002. 10.1016/j.xgen.2021.100002

Frankham, R., Brook, B. W., & Bradshaw, C. J. A. (2014). Genetics in conservation management: revised recommendations for the 50/500 rules, Red List criteria and population viability analyses. Biological Conservation, 170, 56–63. 10.1016/j.biocon.2013.12.036

Franklin, I. R. (1980). Evolutionary change in small populations. Conservation Biology, 135–149. https://cir.nii.ac.jp/crid/1572824500542492672.bib?lang=en

Frichot, E., & François, O. (2015). LEA: An R package for landscape and ecological association studies. Methods in Ecology and Evolution, 6(8), 925–929. 10.1111/2041-210X.12382

Garringer, H. J., Malekpour, M., Esteghamat, F., Mortazavi, S. M. J., Davis, S. I., Farrow, E. G., Yu, X., Arking, D. E., Dietz, H. C., & White, K. E. (2008). Molecular genetic and biochemical analyses of FGF23 mutations in familial tumoral calcinosis. American Journal of Physiology-Endocrinology and Metabolism, 295(4), E929–E937. 10.1152/ajpendo.90456.2008

Glémin, S. (2003). How are deleterious mutations purged? Drift versus nonrandom mating. Evolution, 57(12), 2678–2687. 10.1111/j.0014-3820.2003.tb01512.x

Grossen, C., Guillaume, F., Keller, L. F., & Croll, D. (2020). Purging of highly deleterious mutations through severe bottlenecks in Alpine ibex. Nature Communications, 11(1), 1001. 10.1038/s41467-020-14803-1

Hasselgren, M., Dussex, N., Seth, J., Angerbjörn, A., Dalén, L., & Norén, K. (2024). Strongly deleterious mutations influence reproductive output and longevity in an endangered population. Nature Communications, 15(1), 8378. 10.1038/s41467-024-52741-4

Hedrick, P. W., & Garcia-Dorado, A. (2016). Understanding Inbreeding Depression, Purging, and Genetic Rescue. Trends in Ecology & Evolution, 31(12), 940–952. 10.1016/j.tree.2016.09.005

Hoban, S., Bruford, M., D’Urban Jackson, J., Lopes-Fernandes, M., Heuertz, M., Hohenlohe, P. A., Paz-Vinas, I., Sjögren-Gulve, P., Segelbacher, G., Vernesi, C., Aitken, S., Bertola, L. D., Bloomer, P., Breed, M., Rodríguez-Correa, H., Funk, W. C., Grueber, C. E., Hunter, M. E., Jaffe, R., … Laikre, L. (2020). Genetic diversity targets and indicators in the CBD post-2020 Global Biodiversity Framework must be improved. Biological Conservation, 248, 108654. 10.1016/j.biocon.2020.108654

Huisman, J., Kruuk, L. E. B., Ellis, P. A., Clutton-Brock, T., & Pemberton, J. M. (2016). Inbreeding depression across the lifespan in a wild mammal population. Proceedings of the National Academy of Sciences, 113(13), 3585–3590. 10.1073/pnas.1518046113

Johnston, S. E., Gratten, J., Berenos, C., Pilkington, J. G., Clutton-Brock, T. H., Pemberton, J. M., & Slate, J. (2013). Life history trade-offs at a single locus maintain sexually selected genetic variation. Nature, 502(7469), 93–95. 10.1038/nature12489

Kardos, M., Keller, L. F., & Funk, W. C. (2024). What can genome sequence data reveal about population viability? Molecular Ecology, 0, e17608. 10.1111/mec.17608

Kardos, M., Zhang, Y., Parsons, K. M. A Y., Kang, H., Xu, X., Liu, X., Matkin, C. O., Zhang, P., Ward, E. J., Hanson, M. B., Emmons, C., Ford, M. J., Fan, G., & Li, S. (2023). Inbreeding depression explains killer whale population dynamics. Nature Ecology & Evolution, 7(5), 675–686. 10.1038/s41559-023-01995-0

Keller, L. F., & Waller, D. M. (2002). Inbreeding effects in wild populations. Trends in Ecology & Evolution, 17(5), 230–241. 10.1016/S0169-5347(02)02489-8

Kiernan, J. A. (2015). Histological and Histochemical Methods, fifth edition (5th ed.). Scion Publishing.

Kirkpatrick, M., & Jarne, P. (2000). The Effects of a Bottleneck on Inbreeding Depression and the Genetic Load. The American Naturalist, 155(2), 154–167. 10.1086/303312

Kivisild, T. (2013). Founder Effect. In Brenner’s Encyclopedia of Genetics (pp. 100–101). Elsevier. 10.1016/B978-0-12-374984-0.00552-0

KORA Foundation, Breitenmoser, C., Vogt, K., von Arx, M., Molinari, A., & Breitenmoser, U. (2022). 50 years of lynx presence in Switzerland. KORA Bericht Nr. 99e, 1–80.

KORA Foundation, Breitenmoser, C., Vogt, K., von Arx, M., Signer, S., Zimmermann, F., & Stauffer, C. (2024). Das Projekt LUNO - Abschlussbericht.

Kunz F., Le Grand L., Ziegler E., Bürki R., & Zimmermann F. (2021). Fang-Wiederfang-Schätzung der Abundanz und Dichte des Luchses im Referenzgebiet Simme-Saane IVa im Winter 2020/21. KORA-Bericht Nr. 103.

Kunz F., Ryser J., Breitenmoser-Würsten C., Breitenmoser U., & Zimmermann F. (2020). Fang-Wiederfang-Schätzung der Abundanz und Dichte des Luchses im Berner Oberland Ost IVb im Winter 2019/20. KORA-Bericht Nr. 94.

Marti, I., & Ryser-Degiorgis, M.-P. (2018). A tooth wear scoring scheme for age estimation of the Eurasian lynx (Lynx lynx) under field conditions. European Journal of Wildlife Research, 64. 10.1007/s10344-018-1198-6

Mueller, S., Prost, S., Anders, O., Breitenmoser, C., Kleven, O., Klinga, P., Konec, M., Kopatz, A., Krojerová-Prokešová, J., Middelhoff, L., Obexer-Ruff, G., Reiners, T., Schmidt, K., Sindičić, M., Skrbinšek, T., Tám, B., Saveljev, A., Naranbaatar, G., & Nowak, Carsten. (2022). Genome-wide diversity loss in reintroduced Eurasian lynx populations urges immediate conservation management. Biological Conservation, 266, 109442. 10.1016/j.biocon.2021.109442

Norén, K., Godoy, E., Dalén, L., Meijer, T., & Angerbjörn, A. (2016). Inbreeding depression in a critically endangered carnivore. Molecular Ecology, 25(14), 3309–3318. 10.1111/mec.13674

Pejaver, V., Urresti, J., Lugo-Martinez, J., Pagel, K. A., Lin, G. N., Nam, H.-J., Mort, M., Cooper, D. N., Sebat, J., Iakoucheva, L. M., Mooney, S. D., & Radivojac, P. (2020). Inferring the molecular and phenotypic impact of amino acid variants with MutPred2. Nature Communications, 11(1), 5918. 10.1038/s41467-020-19669-x

Pérez-Pereira, N., Caballero, A., & García-Dorado, A. (2022). Reviewing the consequences of genetic purging on the success of rescue programs. Conservation Genetics, 23(1), 1–17. 10.1007/s10592-021-01405-7

Ralls, K., Ballou, J. D., Rideout, B. A., & Frankham, R. (2000). Genetic management of chondrodystrophy in California condors. Animal Conservation, 3(2), 145–153. 10.1111/j.1469-1795.2000.tb00239.x

Reid, J. M., Arcese, P., & Keller, L. F. (2003). Inbreeding depresses immune response in song sparrows (Melospiza melodia): direct and inter–generational effects. Proceedings of the Royal Society of London. Series B: Biological Sciences, 270(1529), 2151–2157. 10.1098/rspb.2003.2480

Robinson, J. A., Räikkönen, J., Vucetich, L. M., Vucetich, J. A., Peterson, R. O., Lohmueller, K. E., & Wayne, R. K. (2019). Genomic signatures of extensive inbreeding in Isle Royale wolves, a population on the threshold of extinction. Science Advances, 5(5). 10.1126/sciadv.aau0757

Robinson, J., Kyriazis, C. C., Yuan, S. C., & Lohmueller, K. E. (2023). Deleterious Variation in Natural Populations and Implications for Conservation Genetics. Annual Review of Animal Biosciences, 11(1), 93–114. 10.1146/annurev-animal-080522-093311

Roelke, M. E., Martenson, J. S., & O’Brien, S. J. (1993). The consequences of demographic reduction and genetic depletion in the endangered Florida panther. Current Biology, 3(6), 340–350. 10.1016/0960-9822(93)90197-V

Ryser-Degiorgis, M.-P., Robert, N., Meier, R. K., Zürcher-Giovannini, S., Pewsner, M., Ryser, A., Breitenmoser, U., Kovacevic, A., & Origgi, F. C. (2020). Cardiomyopathy associated with coronary arteriosclerosis in free-ranging eurasian lynx (Lynx lynx carpathicus). Frontiers in Veterinary Science, 7. 10.3389/fvets.2020.594952

Sagar, V., Kaelin, C. B., Natesh, M., Reddy, P. A., Mohapatra, R. K., Chhattani, H., Thatte, P., Vaidyanathan, S., Biswas, S., Bhatt, S., Paul, S., Jhala, Y. V., Verma, M. M., Pandav, B., Mondol, S., Barsh, G. S., Swain, D., & Ramakrishnan, U. (2021). High frequency of an otherwise rare phenotype in a small and isolated tiger population. Proceedings of the National Academy of Sciences, 118(39). 10.1073/pnas.2025273118

Sibly, R. M., & Hone, J. (2003). Population growth rate and its determinants: an overview. In R. M. Sibly, J. Hone, & T. H. Clutton-Brock (Eds.), Wildlife Population Growth Rates (pp. 11–40). Cambridge University Press. 10.1098/rstb.2002.1117

Sterrer U., Le Grand L., Kunz F., Rüegg M., & Zimmermann F. (2022). Fang-Wiederfang-Schätzung der Abundanz und Dichte des Luchses in der Nordostschweiz II im Winter 2021/22 KORA-Bericht 109.

Sterrer U., Le Grand L., Künzi E., Lieuwen T., Wolf L., & Zimmermann F. (2025). Fang-Wiederfang-Schätzung der Abundanz und Dichte des Luchses im Referenzgebiet Nordostschweiz II im Winter 2024/2025.

Stoffel, M. A., Johnston, S. E., Pilkington, J. G., & Pemberton, J. M. (2021a). Genetic architecture and lifetime dynamics of inbreeding depression in a wild mammal. Nature Communications, 12(1), 2972. 10.1038/s41467-021-23222-9

Stoffel, M. A., Johnston, S. E., Pilkington, J. G., & Pemberton, J. M. (2021b). Mutation load decreases with haplotype age in wild Soay sheep. Evolution Letters, 5(3), 187–195. 10.1002/evl3.229

Teufel, F., Almagro Armenteros, J. J., Johansen, A. R., Gíslason, M. H., Pihl, S. I., Tsirigos, K. D., Winther, O., Brunak, S., von Heijne, G., & Nielsen, H. (2022). SignalP 6.0 predicts all five types of signal peptides using protein language models. Nature Biotechnology, 40(7), 1023–1025. 10.1038/s41587-021-01156-3

Vogt, K., Korner-Nievergelt, F., Signer, S., Zimmermann, F., Marti, I., Ryser, A., Molinari-Jobin, A., Breitenmoser, U., & Breitenmoser-Würsten, Ch. (2025). Long-Term Changes in Survival of Eurasian Lynx in Three Reintroduced Populations in Switzerland. Ecology and Evolution, 15(4). 10.1002/ece3.71095

Weeks, A. R., Sgro, C. M., Young, A. G., Frankham, R., Mitchell, N. J., Miller, K. A., Byrne, M., Coates, D. J., Eldridge, M. D. B., Sunnucks, P., Breed, M. F., James, E. A., & Hoffmann, A. A. (2011). Assessing the benefits and risks of translocations in changing environments: a genetic perspective. Evolutionary Applications, 4(6), 709–725. 10.1111/j.1752-4571.2011.00192.x

Xue, Y., Prado-Martinez, J., Sudmant, P. H., Narasimhan, V., Ayub, Q., Szpak, M., Frandsen, P., Chen, Y., Yngvadottir, B., Cooper, D. N., de Manuel, M., Hernandez-Rodriguez, J., Lobon, I., Siegismund, H. R., Pagani, L., Quail, M. A., Hvilsom, C., Mudakikwa, A., Eichler, E. E., … Scally, A. (2015). Mountain gorilla genomes reveal the impact of long-term population decline and inbreeding. Science, 348(6231), 242–245. 10.1126/science.aaa3952

