## Supplementary Material for "Evolutionary dynamics of a lethal recessive allele in reintroduced fragmented lynx populations"

#### Supplementary Notes

##### Supplementary Note S1: Histological Evaluation Reveals Multi-Organ Mineralization in Multiple Individuals

Histological re-evaluation revealed nine cases with very similar histopathological findings. All of them showed multifocal to diffuse, moderate to severe tissue mineralization in the kidneys, lung and stomach. In the kidneys most mineral deposits were located in the basement membranes and lumina of both proximal and distal renal tubules. Occasional glomerular tufts showed mineralization, too. Evidence of chronic tissue response, such as fibrosis (i.e. scarring) was observed in five cases. In no case inflammation in the renal tissue was detected in association with the mineralized structures. Pulmonary mineralization predominantly involved the alveolar walls as well as the smooth muscle layer of the bronchi and bronchioles. Associated mild fibrosis was present in half of the cases (n = 5). Of those, two animals exhibited mild inflammatory reactions around mineralized structures. These two, along with four others, also had pneumonia of varying types, not directly related to the mineralization. Across all affected organs, blood vessels consistently showed mineralization of both the lumen and throughout the wall. Furthermore, in the stomach blood vessels represented the most severely affected structure. The extent of mineralization within the different stomach layers varied among cases, though the tunica mucosa was most frequently affected.

More details on pathological findings and sample backgrounds are compiled in *Supplementary Tables S1 and S2* (provided as an Excel file).

##### Supplementary Note S2: Pathogenicity Predictions of the XP_046921148.1:p.Leu13Pro Variant

The reference sequences of the FGF23 protein precursors are 100% identical between bobcat (*Lynx rufus*, XP_046921148.1), Canadian lynx (*Lynx canadiensis*, XP_030177644.1), and domestic cat (*Felis catus*, XP_011282059.2), and consist of 245 amino acids. Based on our WGS data, the FGF23 precursor protein from healthy Eurasian lynx (*Lynx lynx*) shares the same amino acid sequence. We therefore used the *L. rufus* accession XP_046921148.1 for all pathogenicity analyses of the L13P missense variant (Suppl. Figure S5).

We ran three different pathogenicity prediction tools to evaluate the effect of the L13P variant. SNPs&GO predicted a disease-causing effect with a probability of 73.4% with moderate reliability index (RI=5). The PhD-SNP method that is part of SNPs&GO predicted a disease probability of 92.9% with high confidence (RI=9; (Calabrese et al., 2009). Polyphen-2 predicted the variant to be probably damaging with a score of 1.000 in the HumDiv and a score of 0.972 in the HumVar setting (Adzhubei et al., 2013). The MutPred2 analysis resulted in a general score of 0.819, indicating a strong likelihood of pathogenicity (Pejaver et al., 2020). Predicted impact on the molecular mechanisms included loss of helix with a 28% chance for structural change to occur and a possible altered signal peptide with a lower probability of 6%.

In order to more specifically investigate the effect of the L13P variant on signal peptide recognition, we compared the wildtype and mutant sequence with the SignalP 6.0 software (Teufel et al., 2022). The FGF23 signal peptide belongs to the canonical Sec/SPI class of mammalian signal peptides that are characterized by a positively charged N-region of 1-5 aa, a hydrophobic α-helix consisting of 7-15 aa, which is termed H-region and a C-terminal C-region of 3-7 aa. The central hydrophobic α-helix assists with targeting, embedding, and translocating the nascent peptide across the cell membrane during the transport process (Zhang et al., 2025). The SignalP 6.0 software recognized the signal peptide and cleavage site in the wildtype FGF23 sequence with a probability of 98.2%, while this value dropped to 77.7% in the L13P mutant sequence. Proline is a known breaker of α-helical secondary structures. Leucine to proline substitutions in the H-region of signal peptides have been observed in human inherited diseases. An L6P substitution in the human FGF3 protein represents a known pathogenic variant and leads to congenital deafness with inner ear agenesis, microtia and microdontia (Tekin et al., 2008). Taken together, the *in silico* predicted effects of the lynx FGF23:p.L13P variant suggest that the mutant allele may reduce or even abolish the correct posttranslational sorting, cleavage and secretion of FGF23 and are compatible with a causal role of this variant for the observed mineralizations in lynx.

#### Supplementary Tables

**Supplementary Table S1** (provided in Excel file Supplementary_Tables_S1&S2.xlsx): Pathological description of the nine cases.

**Supplementary Table S2** (provided in Excel file Supplementary_Tables_S1&S2.xlsx): Sample info of whole-genome sequencing and targeted genotyping.

**Supplementary Table S3**. Variant Filtering Steps and Subsequent Analyses with resulting Variant Counts

| Filtering Step | Variant Count |
|---|---|
| Raw vcf after GATK Variant Filtration | 17,328,023 |
| Bi-allelic only, QD>2, minGQ 20, max-missing 0.8, maf 0.05, autosomes only | 1,899,640 |
| QD>15 | 1,785,573 |
| GEMMA | 1,699,701 |

**Supplementary Table S4.** Primer Sequences and PCR Conditions

| | Forward Primer | Reverse Primer |
|---|---|---|
| **Name** | Lynx_FGF23_F | Lynx_FGF23_R |
| **Gene** | FGF23 | FGF23 |
| **Chromosome** | B4 | B4 |
| **Position start (NW_025814889.1)** | 39,312,463 | 39,312,691 |
| **Position stop (NW_025814889.1)** | 39,312,482 | 39,312,672 |
| **Sequence** | ATGGCCGTCCTTGTGTATCT | AGACCAGGGAGTCAGGGAGT |
| **Length [bp]** | 20 | 20 |
| **Tm [°C]** | 59.4 | 60.1 |
| **GC [%]** | 50 | 60 |
| **Product Length [bp]** | 229 | 229 |
| **Comments** | 1 mismatch to *Lynx rufus* genome, perfect match to *Lynx lynx* sequences (*Lynx rufus*:ATGGCCATCCTTGTGTATCT) | |

#### Supplementary Figures

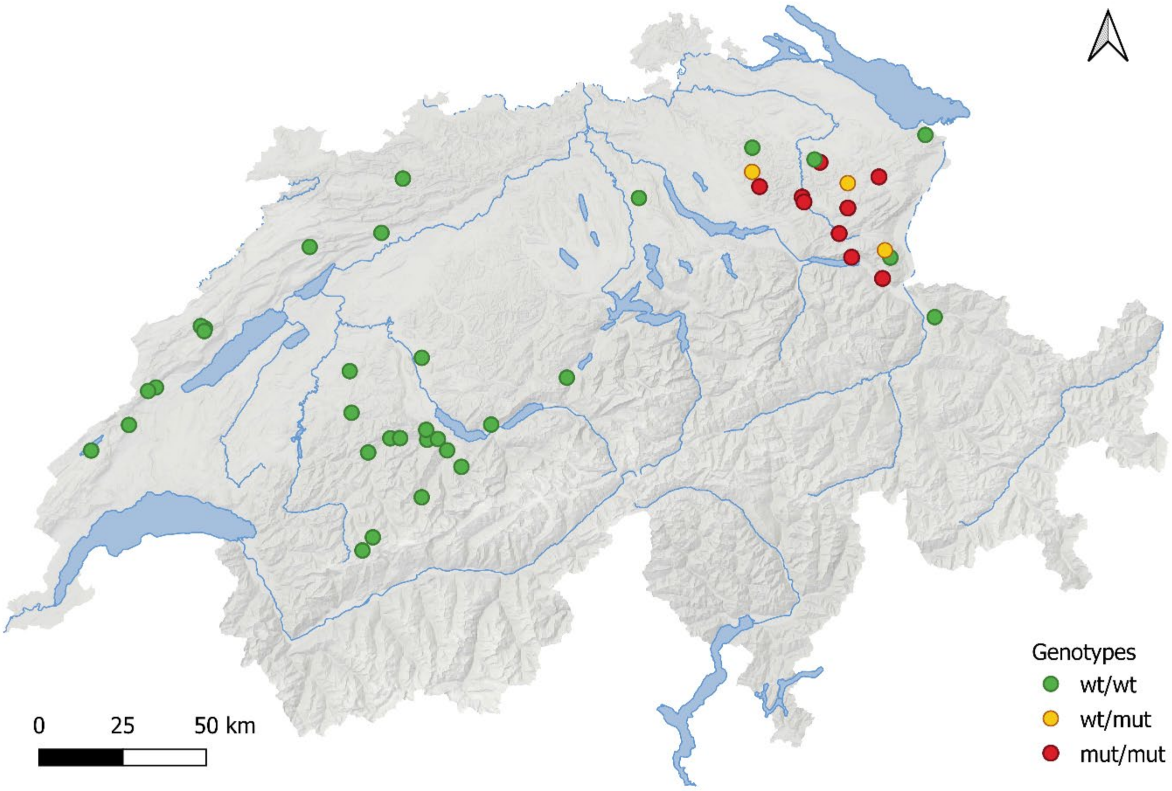


**Supplementary Figure S1**. Map of Switzerland showing lynx included in the whole-genome sequencing (WGS). Dots indicate locations where animals were found dead or sampled during live captures, color-coded by genotype. Red dots indicate affected individuals homozygous for the lethal allele.

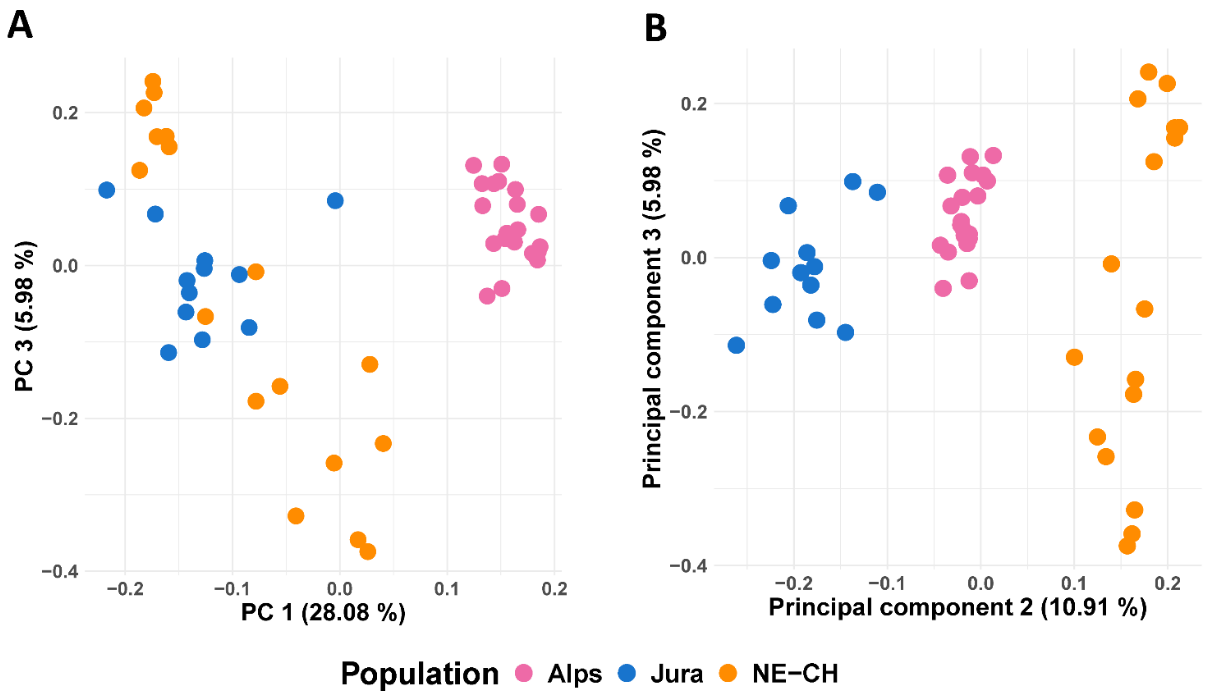


**Supplementary Figure S2**. Principal Component Analysis (PCA) revealed clear genetic structure among the three populations. Principal component 1 (PC1, 28.08 % of the variance) separated the Alpine population from both Jura and NE-CH, but not the latter two from each other (A), consistent with a larger proportion of Jura individuals having contributed to the NE-CH population (KORA, 2024). PC2 (10.91%) further differentiated all three populations, particularly separating NE-CH from Jura (B). PC3 accounted for only 5.98% of the total variance, and individuals from all three populations overlapped along this axis, indicating no clear genetic differentiation along PC3.

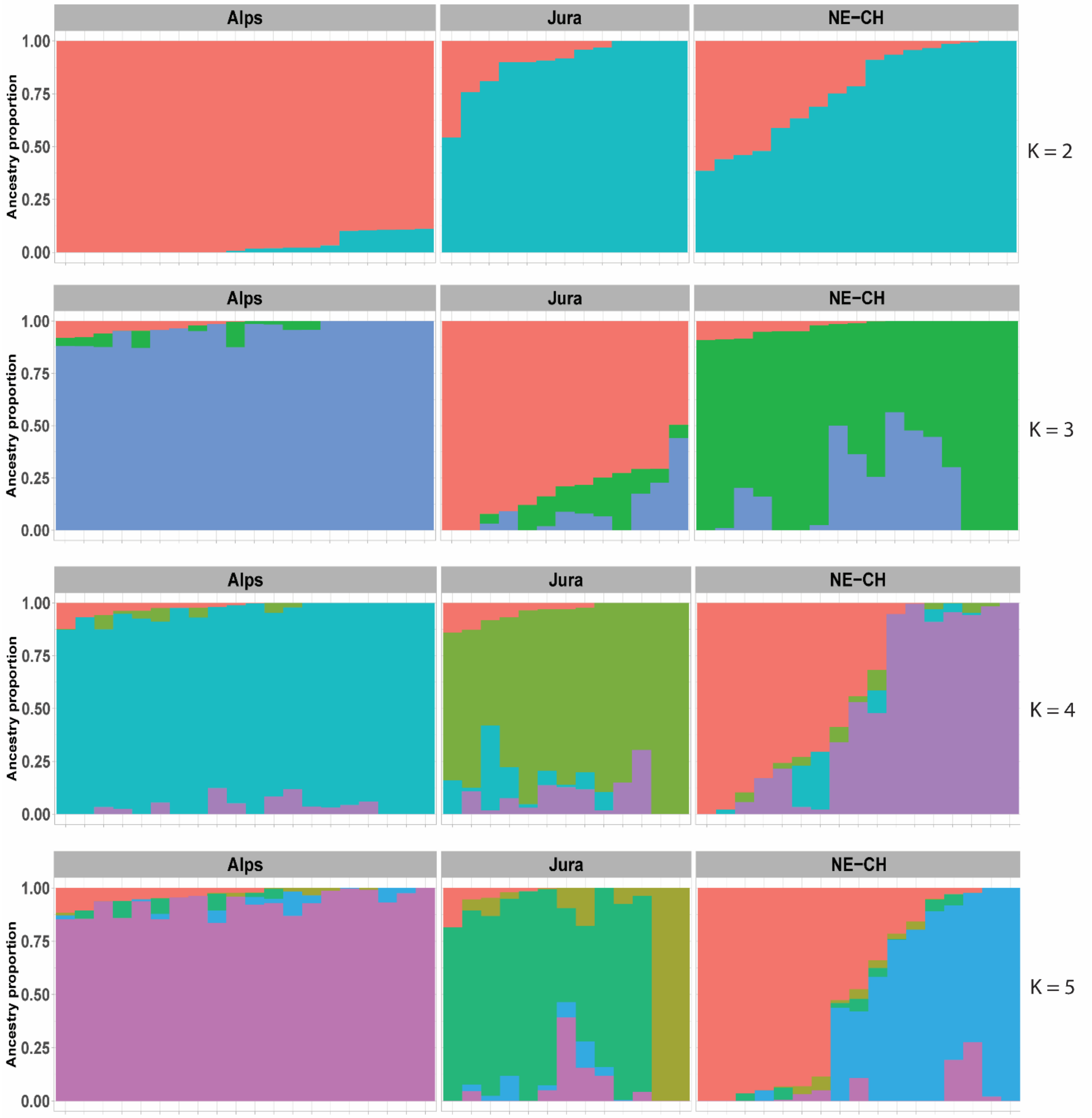


**Supplementary Figure S3** Population structure estimated using the sNMF clustering algorithm, testing K = 2–5 ancestral populations. Results generally confirm the genetic separations suggested by the PCA. At K = 2 individuals from the Alps and Jura form two genetic clusters, with a few individuals of mixed ancestry. At K = 3, NE-CH individuals form a separate cluster, reflecting their genetic differentiation from the other two populations.

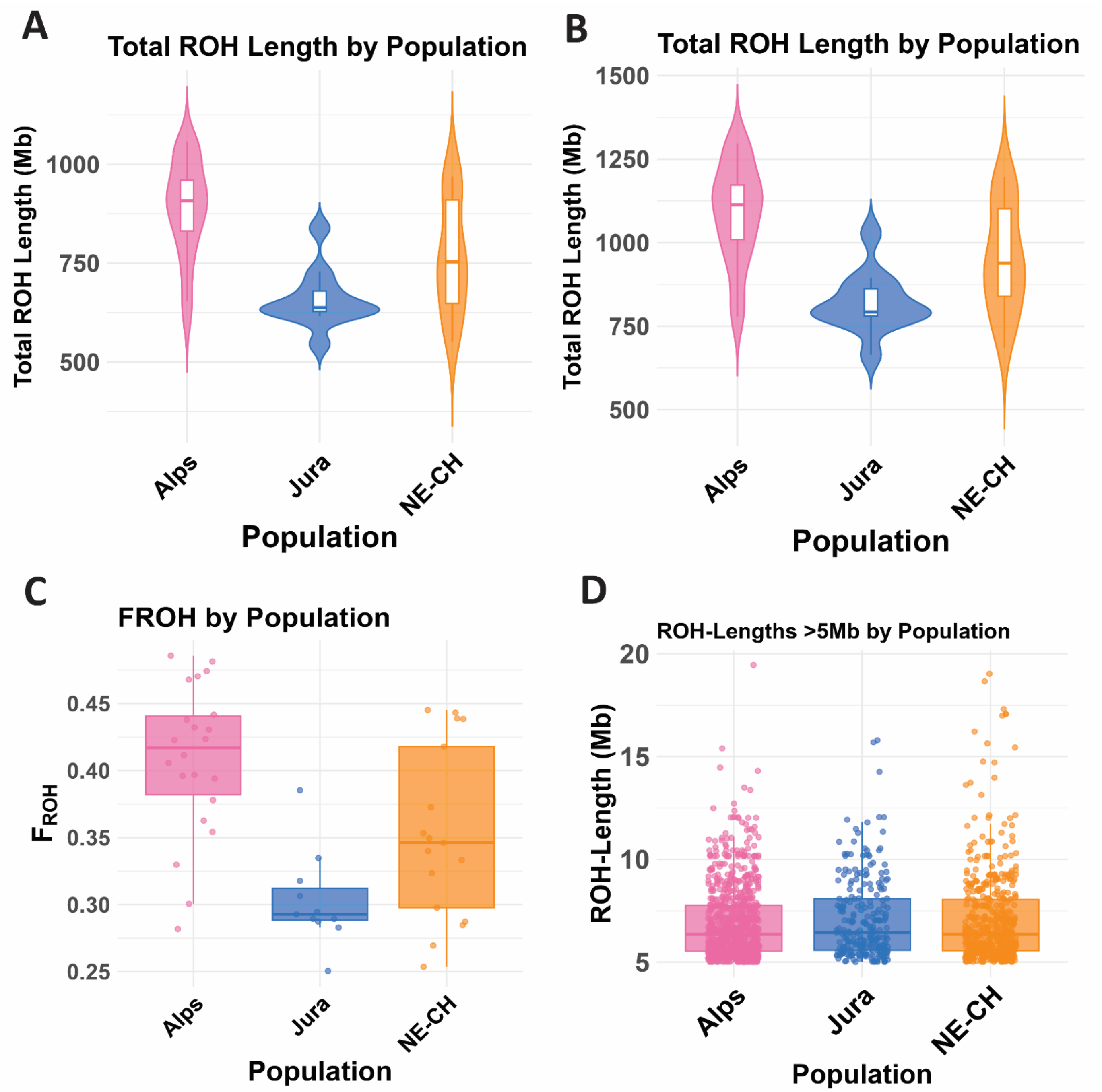


**Supplementary Figure S4**. Total ROH Length Analysis of ROH > 1 Mb with QD>2 (A) and QD>15 (B). FROH with QD>2 (C) and ROH-length distribution for ROH >5Mb using the dataset filtered with QD>15. ROH analyses based on less stringently filtered variants (QD > 2) let to slightly lower sROH and FROH values, but overall qualitative patterns across populations remained consistent. When comparing ROH lengths >5 Mb and >10 Mb, the majority of ROH were shorter than 10 Mb.

**A**

L13P ↓

```
  1 MSGTRLGLLV SVLCWVVRAY PNTSPLLGSS WGGLTHLYTA TARNSYHLQI HKDGHVDGTP
 61 HQTIYSALMI RSEDAGFVVI TGVMSQRYLC MDFRGNIFGS HLFSPESCRF RQRTLENGYD
121 VYHSPQHRFL VSLGPAKRAF LPGTNPPPYS QFLSRRNEIP LVHFNTPRPR RHTRSAEDAE
181 RDPLNVLKPR PRMTPAPASC SQELPSAEDS GVVASDPLGV LRGNRVNAHA GGMGVERCRP
241 FPKFN
```

**B** SignalP 6.0 wildtype

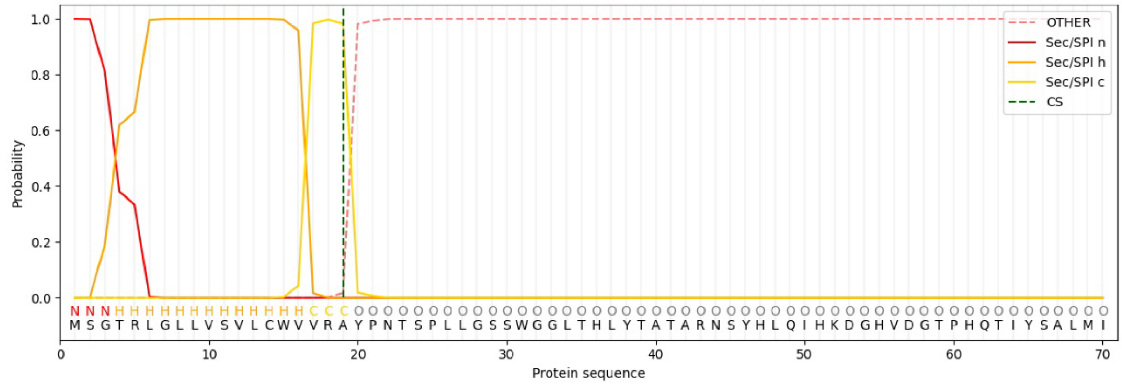


**C** SignalP 6.0 L13P mutant

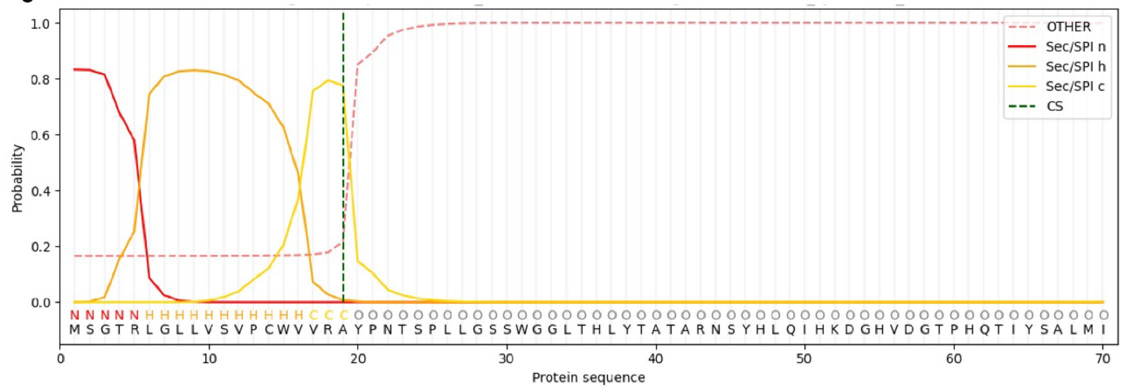


**Supplementary Figure S5**. FGF23 protein analyses. (A) Amino acid sequence of the FGF23 precursor protein from *L. rufus*, accession XP_046921148.1. The signal peptide comprises amino acids 1 – 19 and is highlighted in orange. (B, C) Signal peptide prediction with SignalP 6.0 of the wildtype and L13P mutant FGF23 sequence. The probability for signal peptide recognition decreases in the mutant sequence.

### References


Adzhubei, I., Jordan, D. M., & Sunyaev, S. R. (2013). Predicting functional effect of human missense mutations using PolyPhen-2. *Current Protocols in Human Genetics*, *76*(1). https://doi.org/10.1002/0471142905.hg0720s76

Calabrese, R., Capriotti, E., Fariselli, P., Martelli, P. L., & Casadio, R. (2009). Functional annotations improve the predictive score of human disease-related mutations in proteins. *Human Mutation*, *30*(8), 1237–1244. https://doi.org/10.1002/humu.21047

KORA Foundation, Breitenmoser, C., Vogt, K., von Arx, M., Signer, S., Zimmermann, F., & Stauffer, C. (2024). *Das Projekt LUNO - Abschlussbericht*.

Pejaver, V., Urresti, J., Lugo-Martinez, J., Pagel, K. A., Lin, G. N., Nam, H.-J., Mort, M., Cooper, D. N., Sebat, J., Iakoucheva, L. M., Mooney, S. D., & Radivojac, P. (2020). Inferring the molecular and phenotypic impact of amino acid variants with MutPred2. *Nature Communications*, *11*(1), 5918. https://doi.org/10.1038/s41467-020-19669-x

Tekin, M., Öztürkmen Akay, H., Fitoz, S., Birnbaum, S., Cengiz, F., Sennaroğlu, L., İncesulu, A., Yüksel Konuk, E., Hasanefendioğlu Bayrak, A., Şentürk, S., Cebeci, İ., Ütine, G., Tunçbilek, E., Nance, W., & Duman, D. (2008). Homozygous *FGF3* mutations result in congenital deafness with inner ear agenesis, microtia, and microdontia. *Clinical Genetics*, *73*(6), 554–565. https://doi.org/10.1111/j.1399-0004.2008.01004.x

Teufel, F., Almagro Armenteros, J. J., Johansen, A. R., Gíslason, M. H., Pihl, S. I., Tsirigos, K. D., Winther, O., Brunak, S., von Heijne, G., & Nielsen, H. (2022). SignalP 6.0 predicts all five types of signal peptides using protein language models. *Nature Biotechnology*, *40*(7), 1023–1025. https://doi.org/10.1038/s41587-021-01156-3

Zhang, S., He, Z., Wang, H., & Zhai, J. (2025). Signal Peptides: From molecular mechanisms to applications in protein and vaccine engineering. *Biomolecules*, *15*(6), 897. https://doi.org/10.3390/biom15060897
